# FOXA2 is essential for maintaining the urea cycle in acute liver failure

**DOI:** 10.1101/2023.12.11.570955

**Authors:** Rilu Feng, Rui Liu, Chenhao Tong, Tao Lin, Xiaofeng Li, Hui Liu, Chen Shao, Kejia Kan, Carsten Sticht, Yujia Li, Sai Wang, Stefan Munker, Ulrich Wirth, Hanno Niess, Roman Liebe, Christoph Meyer, Shanshan Wang, Matthias P Ebert, Steven Dooley, Huiguo Ding, Hua Wang, Hong-Lei Weng

**Author notes:** Corresponding author: Hua Wang. Hong-Lei Weng. The authors contributed equally and share first authorship (Rilu Feng, Rui Liu). These authors contributed equally and share corresponding authorship (Hua Wang, Hong-Lei Weng).

## Abstract

Hepatic encephalopathy is a lethal complication of acute liver failure (ALF), and is caused by hyperammonemia. Ammonia clearance by the liver requires an intact and complete urea cycle comprising six enzymes, including the rate-limiting enzyme carbamoyl phosphate synthetase I (CPS1). To date, the detailed regulation of CPS1 transcription in order to maintain urea cycle in physiological condition and ALF remains largely unknown. This study scrutinizes the role of pioneer factor forkhead box A 2 (FOXA2) in the regulation of CPS1 transcription, urea cycle performance and hyperammonemia. Physiologically, CPS1 transcription requires FOXA2 to maintain chromatin accessibility on its enhancers, which is essential for CCAAT enhancer-binding protein-alpha (C/EBPα) binding to activate gene transcription. In ALF, hepatic C/EBPα expression is inhibited by inflammatory mediators such as TGF-β and TNF-α. In this setting, retinoic acid receptor synergizes with FOXA2 to maintain CPS1 transcriptions. Once ALF patients suffer from massive hepatic necrosis, liver progenitor cells initiate a transcription network comprising FOXA2 and C/EBPα to perform the urea cycle and prevent hyperammonemia. In ALF, hepatic encephalopathy occurs in patients lacking hepatic FOXA2 expression. In mice with acetaminophen-induced ALF, injection of Foxa2-AAV8 maintains urea cycle and prevents hyperammonemia. Taken together, FOXA2 is essential for maintaining the urea cycle. Pharmaceutical induction of hepatic FOXA2 expression might represent a novel approach to treat hepatic encephalopathy in ALF.

**One Sentence Summary:** Pioneer factor FOXA2 synergizes with C/EBPα or RAR to maintain urea cycle in acute liver failure

## INTRODUCTION

Acute liver failure (ALF) is a medical emergency characterized by two severe pathophysiological alterations: coagulation abnormality and hepatic encephalopathy (HE) (*1*). In ALF, HE is induced by severe inflammation and hyperammonemia, which is caused by massive hepatocyte loss and impaired urea cycle in remaining hepatocytes (*2*). The severity of HE is closely associated with prognosis of ALF (*2*).

The urea cycle is mainly performed by hepatocytes. A complete cycle comprises six enzymes: carbamoyl phosphate synthase I (CPS1), ornithine transcarbamylase (OTC) and N-acetylglutamate synthase (NAGS) in mitochondria and argininosuccinate synthase (ASS), argininosuccinate lyase (ASL) and arginase 1 (ARG1) in cytoplasm (*3*). Among these enzymes, CPS1 catalyzes the irreversible first step of the cycle and plays a rate-limiting role (*3*). CPS1 transcription is regulated by multiple transcription factors, including CCAAT enhancer-binding protein-alpha (C/EBPα), which possesses binding sequences on the promoter and enhancer of the CPS1 gene (*4*). Cebpa-deficient mice suffer from hyperammonemia 5. In ALF, C/EBPα expression in hepatocytes is often inhibited by inflammatory factors (*6*). Furthermore, severe ALF patients suffer from massive hepatocyte loss (*7*). However, a portion of ALF patients with massive hepatocyte loss lacking C/EBPα expression do not develop hyperammonemia and HE, indicating the existence of a C/EBPα- and hepatocyte-independent rescue network for the maintenance of urea cycle to prevent hyperammonemia in ALF.

This study outlines a hierarchical network that maintains urea cycle in health and ALF: (1) Physiologically, CPS1 transcription requires a pioneer factor forkhead box A2 (FOXA2) to maintain chromatin accessibility on its enhancer, which is essential for C/EBPα binding to activate gene transcription. (2) When hepatic C/EBPα expression is inhibited by inflammation, retinoic acid receptor (RAR) deputies C/EBPα and collaborates with FOXA2 to maintain CPS1 transcription. (3) Once ALF patients suffer from massive hepatic necrosis, liver progenitor cells (LPCs) perform the urea cycle to prevent hyperammonemia by initiating a transcription network comprising FOXA2 and C/EBPα.

## RESULTS

### Patients with hepatic encephalopathy lack FOXA2 expression

First, immunohistochemical staining (IHC) was performed to examine expression of CPS1 in the collected liver tissues. Expression of CPS1 was only detectable in patients without HE, but not in those with HE (**Figure 1**). In ALF patients without HE, CPS1 was expressed in both hepatocytes and LPCs (**Figure 1**). Impressively, IHC further revealed that the hepatocytes and LPCs expressing CPS1 also presented robust FOXA2 expression (**Figure 1 and Table S1**). These results suggest a potential relationship between FOXA2 and CPS1 in the occurrence of HE.

**Fig. 1.**
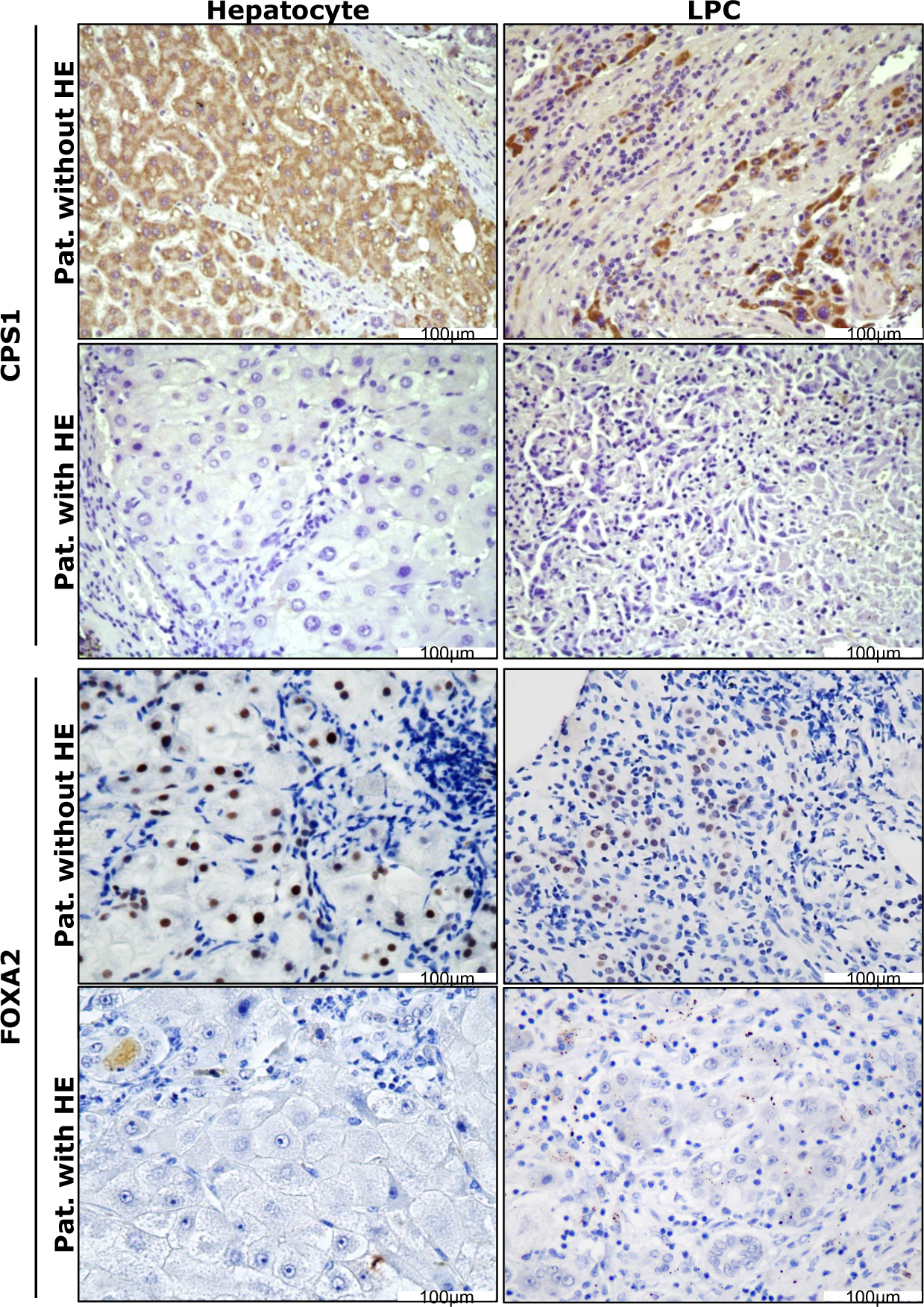
Hepatic CPS1 and FOXA2 expression in patients with and without hepatic encephalopathy. Immunohistochemical staining for FOXA2, and CPS1 was performed in patients with acute liver failure. Representative patients show that CPS1 and FOXA2 are expressed in hepatocytes (left panel) and liver progenitor cells (LPC, right panel) in patients without hepatic encephalopathy (HE), but not in those suffering from HE. Original magnification: x20.

### FOXA2 regulates urea cycle through regulating *CPS1* gene

To clarify the effects of FOXA2 on urea and ammonia products in hepatocytes, *Foxa2* construct or siRNA were transfected into the mouse hepatocyte cell line AML12 and mouse primary hepatocytes (MPHs), respectively. Overexpression of *Foxa2* increased urea and reduced ammonia concentrations in supernatants of hepatocytes, whereas knockdown of *Foxa2* resulted in opposite effects (**Figure 2A**). DNA sequence analysis further revealed potential FOXA2 binding sites on the *Cps1* (-718∼-260bp) promoter (**Figure 2B**), which were subsequently confirmed by ChIP assay (**Figure 2C**). ChIP-seq performed in mouse liver further showed that FOXA2 bound to several sites of the *Cps1* gene, including the sites described above (GSE157452, **Figure 2D**). qPCR analyses revealed that knockdown of *FOXA2* by RNAi reduced mRNA expression of CPS1 in AML12 cells, MPHs and human primary hepatocytes (HPHs) (**Figure 2E**). Western blotting performed in AML12 cells and MPHs further showed reduced protein expression of CPS1 when FOXA2 was knocked down (**Figure 2F**). Interestingly, overexpression of *FOXA2* did not remarkably influence mRNA expression of *CPS1* in three types of hepatocytes (**Figure 2G**). In HPH, overexpression of *FOXA2* even slightly reduced *CPS1* expression (**Figure 2G**).

**Fig. 2.**
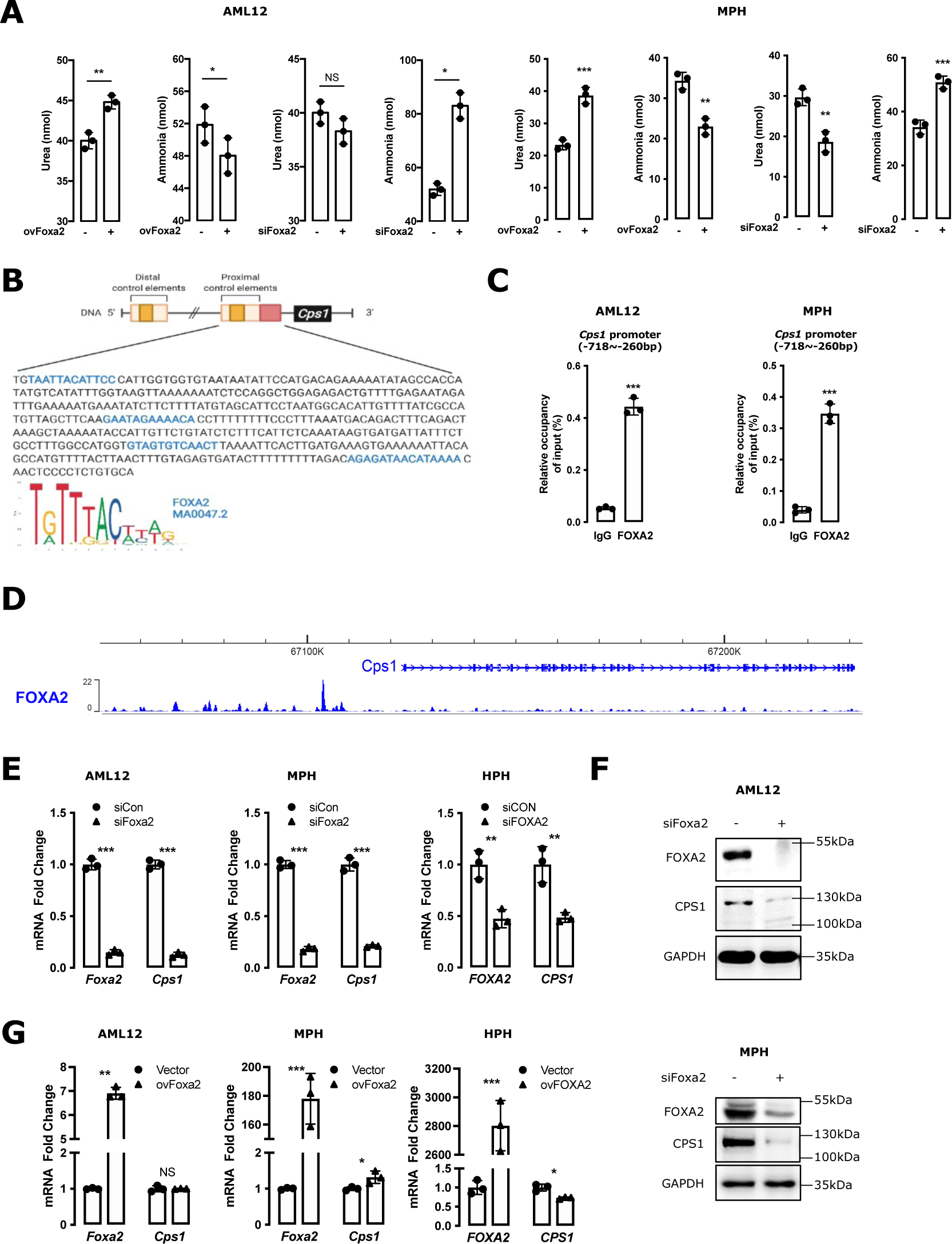
FOXA2 regulates urea cycle through binding to the *CPS1* promoter. **(A)** Urea and ammonia levels were examined in the culture medium of AML12 cells and mouse primary hepatocytes (MPH) transfected with Foxa2 construct or siRNA for 48 hours. **(B)** The binding sites of FOXA2 on *Cps1* (-718∼-260bp) promoter were predicted by Jaspar dataset (*26*). The putative FOXA2 binding sites are dyed blue. **(C)** ChIP assays examined FOXA2 binding to the *Cps1* (-718∼-260bp) promoter in AML12 cells and MPH. **(D)** ChIP sequencing (ChIP-seq) showed enrichment profiles of FOXA2 on *Cps1* promoter in mouse liver tissue. **(E-F)** qPCR **(E)** and western blotting **(F)** analyzed the effects of FOXA2 on CPS1 expression in AML12 cells, MPH, and human primary hepatocytes (HPH). The cells were transfected with FOXA2 siRNA for 48 hours. **(G)** qPCR examined the effects of FOXA2 on CPS1 expression in the cells transfected with FOXA2 construct for 48 hours. Bars represent the mean ± SD, P values were calculated by unpaired Student’s t-test, *, P<0.05; **, P<0.01, and ***, P<0.001 and NS, no significance. Triple experiments were performed, and one representative result is shown.

These results suggest that FOXA2 might be an essential, but not sufficient factor for CPS1 expression in hepatocytes.

### FOXA2 regulates CPS1 transcription as a pioneer factor

FOXA2 usually acts as a pioneer factor in the regulation of gene transcription (*8*). A previous study showed that FOXA2 opens chromatin at the albumin (*ALB*) enhancer, which is essential for subsequent C/EBPα and HNF4α binding to the enhancer to regulate *ALB* transcription (*6*). Notably, overexpression of *FOXA2* exclusively does not significantly increase *ALB* expression (*6*). Therefore, FOXA2 might also function as a pioneer factor in the transcriptional regulation of *CPS1*.

As in *ALB* transcription, C/EBPα has previously been recognized as critical transcription factors in the regulation of CPS1 expression (*4*). In AML12 cells and MPHs, knockdown of *Cebpa* remarkably reduced mRNA expression of Cps1 (**Figure 3A**). In those hepatocytes with *Cebpa* knockdown, overexpression of *Foxa2* did not rescue mRNA expression of Cps1 (**Figure 3A**), indicating an indispensable effect of C/EBPα on FOXA2-regulated CPS1 transcription.

**Fig. 3.**
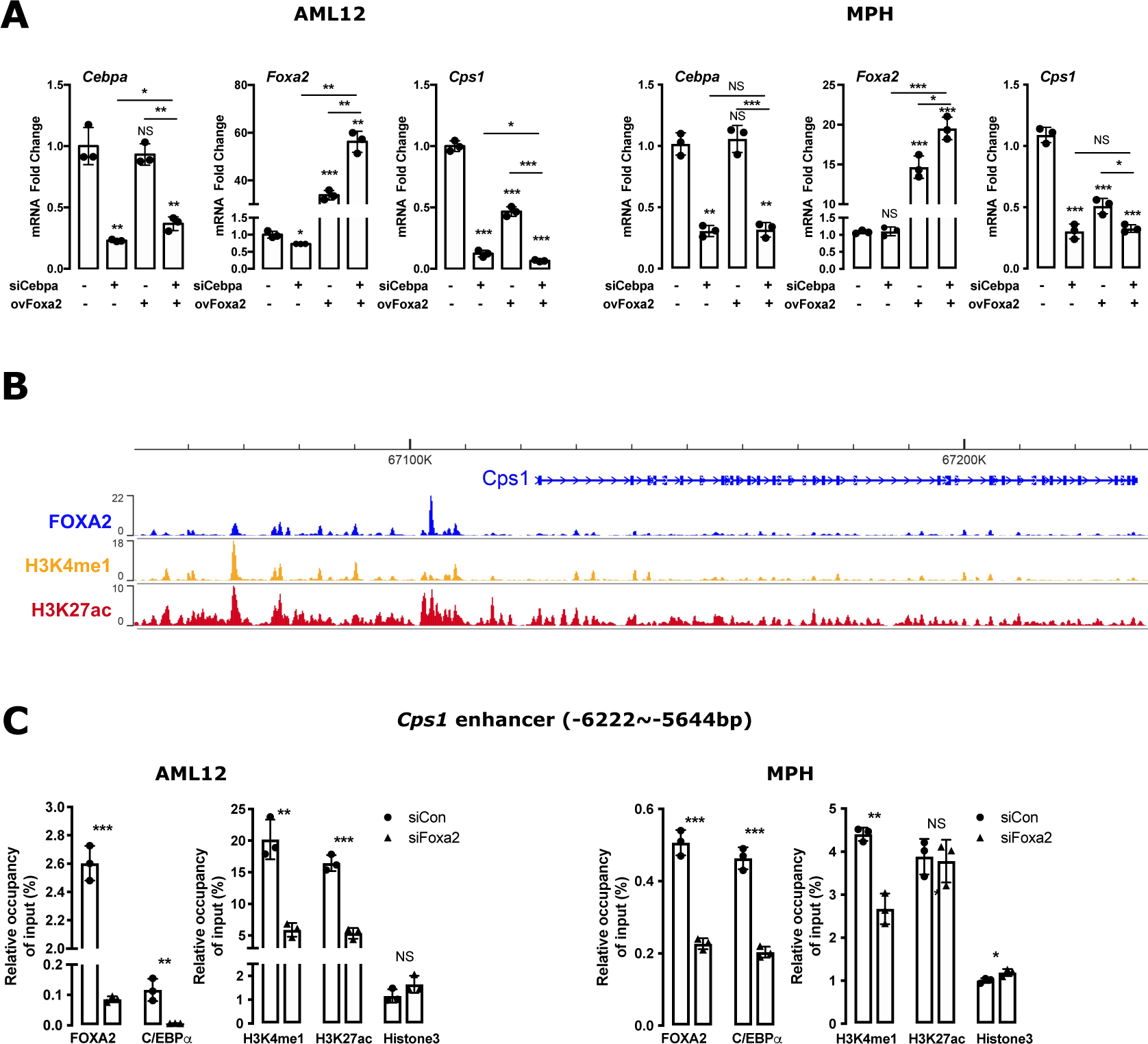
FOXA2 regulates *CPS1* gene as a pioneer factor. **(A)** Following siRNA-Cebpa transfection for 24 hours, AML12 cells and mouse primary hepatocytes (MPH) were transfected by Foxa2 construct for additional 48 hours. qPCR analyzed the effects of FOXA2 on Cps1 expression in both cells with Cebpa disruption. **(B)** The enrichment profiles for FOXA2, H3K4me1 and H3K27ac across a 10 kbp region of the *Cps1* gene in mouse liver tissue. **(C)** ChIP assays were performed to examine FOXA2, C/EBPα, and Histone proteins on *Cps1* (-6222∼-5644bp) enhancer in AML12 cells and MPH with and without Foxa2 knockdown. Bars represent the mean ± SD, P values were calculated by One-way ANOVA, *, P<0.05; **, P<0.01, and ***, P<0.001 and NS, no significance. Triple experiments were performed, and one representative result is shown.

To further clarify the role of FOXA2 in the *CPS1* gene transcription, a ChIP-seq database (GSE157452) performed in mouse livers was analyzed. FOXA2 robustly bound to the *Cps1* enhancer at approximately -6.3 kbp and -7.8 kbp from upstream of the transcription start site (**Figure 3B**), the region also comprising C/EBPα binding sites (*4*). With FOXA2 binding, these regions show high levels of H3K4me1 and H3K27ac, two markers of chromatin accessibility (**Figure 3B**), suggesting a critical role of FOXA2 in maintaining chromatin accessibility to the enhancer.

Subsequently, ChIP assays were performed in AML12 cells and MPHs to examine whether FOXA2 disruption influenced C/EBPα binding to the *CPS1* enhancer. As expected, C/EBPα binding to the *Cps1* enhancer at -6222 ∼ -5644bp was remarkably inhibited in the cells with *Foxa2* knockdown (**Figure 3C**). Furthermore, Histone H3 binding at these regions was significantly increased, whereas H3K4me1 and H3K27ac bindings were reduced (**Figure 3C**). These results suggest that FOXA2 acts as a pioneer factor to maintain chromatin accessibility, which is essential for C/EBPα in the regulation of *CPS1* transcription in hepatocytes.

### C/EBPα regulates *CPS1* transcription in hepatocytes

Given the key role of C/EBPα in the regulation of CPS1 (*4*), RNA sequencing was performed in MPHs with or without *Cebpa* gene knockdown. Multiple arginine biosynthesis relevant transcripts, including Cps1 were altered in MPHs with *Cebpa* knockdown (**Figure 4A-B**). qPCR analysis confirmed that mRNA expression of Cps1, Otc, Nags, and Arg1 was inhibited when *Cebpa* was knocked down by RNAi in AML12 cells (**Figure 4C**).

**Fig. 4.**
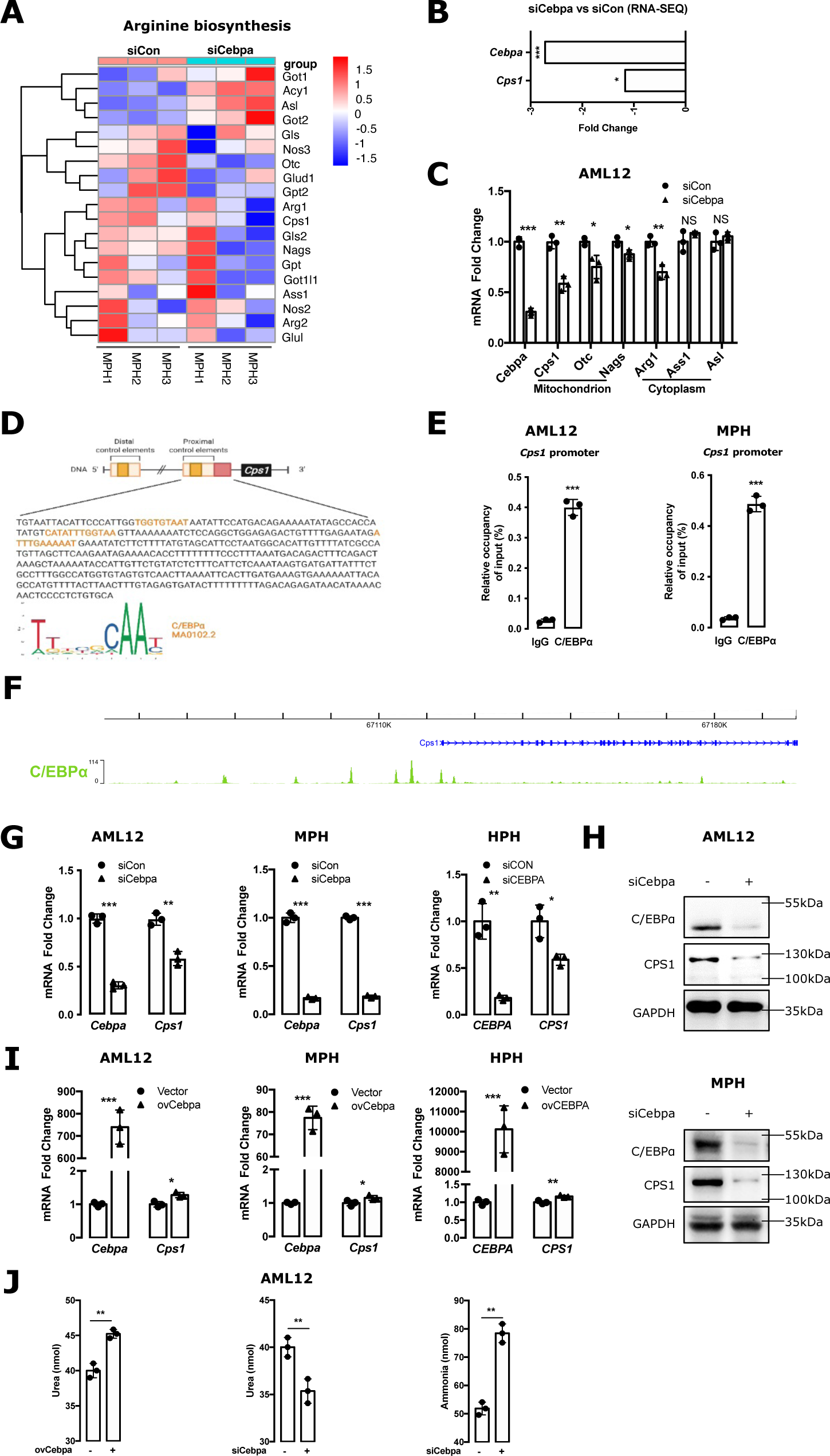
C/EBPα regulates CPS1 transcription in hepatocytes. **(A)** A heatmap depicts expression profile of arginine biosynthesis in mouse primary hepatocytes (MPH) with Cebpa knockdown by RNAi. **(B)** RNA sequencing shows altered expression of Cebpa and Cps1 in MPH with Cebpa knockdown. **(C)** qPCR was performed to analyze the effects of C/EBPα on components of urea cycle in AML12 cells. The cells were transfected with Cebpa siRNA for 48 hours. **(D)** The binding sites of C/EBPα on *Cps1* promoter (-718∼- 260bp) were predicted by Jaspar dataset (*26*). The putative C/EBPα binding sites are dyed orange. **(E)** ChIP assays confirmed the binding of C/EBPα on *Cps1* (-718∼-260bp) promoter in AML12 cells and MPH. **(F)** ChIP sequencing (ChIP-seq) showed enrichment profiles of C/EBPα on *Cps1* promoter in mouse liver tissue. **(G-H)** qPCR **(G)** and western blotting **(H)** analyzed the effects of C/EBPα on CPS1 expression in AML12 cells, MPH, and human primary hepatocytes (HPH). The cells were transfected with CEBPA siRNA for 48 hours. **(I)** qPCR examined the effects of C/EBPα on CPS1 expression in the cells transfected with CEBPA construct for 48 hours. **(J)** Urea and ammonia levels were measured in the culture medium of AML12 cells transfected with Cebpa construct or siRNA for 48 hours. Bars represent the mean ± SD, P values were calculated by unpaired Student’s t-test, *, P<0.05; **, P<0.01, and ***, P<0.001. Triple experiments were performed, and one representative result is shown.

Subsequently, the effect of C/EBPα on CPS1 expression was further investigated. Besides binding to the *Cps1* enhancer (*4*), DNA sequence analysis showed that C/EBPα also bound to the *Cps1* promoter (-718∼-260bp) (**Figure 4D**). ChIP assay confirmed the binding of C/EBPα on the *Cps1* promoter in both AML12 cells and MPHs (**Figure 4E**). ChIP-seq performed in mouse liver further showed that C/EBPα bound to several sites of the *Cps1* gene, including the sites described above (GSE65167, **Figure 4F**). qPCR and western blotting analyses further showed that knockdown of *CEBPA* inhibited CPS1 expression in AML12 cells, MPHs and HPHs (**Figure 4G-H**). Interestingly, overexpression of *CEBPA* only slightly increased mRNA expression of CPS1 in three hepatocytes (**Figure 4I**). Overexpression of *Cebpa* increased urea levels, while knockdown of *Cebpa* reduced urea levels and increased ammonia concentrations in supernatant of the AML12 cells (**Figure 4J**). Furthermore, IHC showed that ALF patients with HE did not have detectable C/EBPα expression (**Table S1**). These results suggest a crucial role of C/EBPα in the regulation of CPS1 expression and urea cycle in hepatocytes.

### RAR is a key factor to regulate urea cycle in hepatocytes lacking C/EBPα

In ALF patients, C/EBPα expression is often undetectable due to severe inflammation (*6*). In a previous study, most irreversible ALF patients regardless of HE did not have detectable expression of C/EBPα (*6*). However, a large portion of patients do not suffer from HE in the absence of C/EBPα. Which transcription factor replaces C/EBPα to regulate urea cycle in these patients? A previous study showed that RAR, supported by FOXA2, replaces C/EBPα and HNF4α to regulate *ALB* transcription (*6*). Therefore, RAR might also act as an alternative factor to regulate transcription of the urea cycle genes.

In AML12 cells and HPHs, administration of all trans retinoic acid (ATRA) time-dependently increased mRNA expression of CPS1 (**Figure 5A**). Furthermore, ATRA still significantly induced CPS1 expression in hepatocytes with *CEBPA* knockdown (**Figure 5B-C**). DNA sequence analyses showed RAR binding sites on the *Cps1* promoter (**Figure 5D**). Subsequent ChIP assay confirmed the binding of RAR on the *Cps1* (-718∼-260bp) promoter in AML12 cells (**Figure 5E**). ATRA stimulation further increased RAR binding on the *Cps1* promoter (**Figure 5E**).

**Fig. 5.**
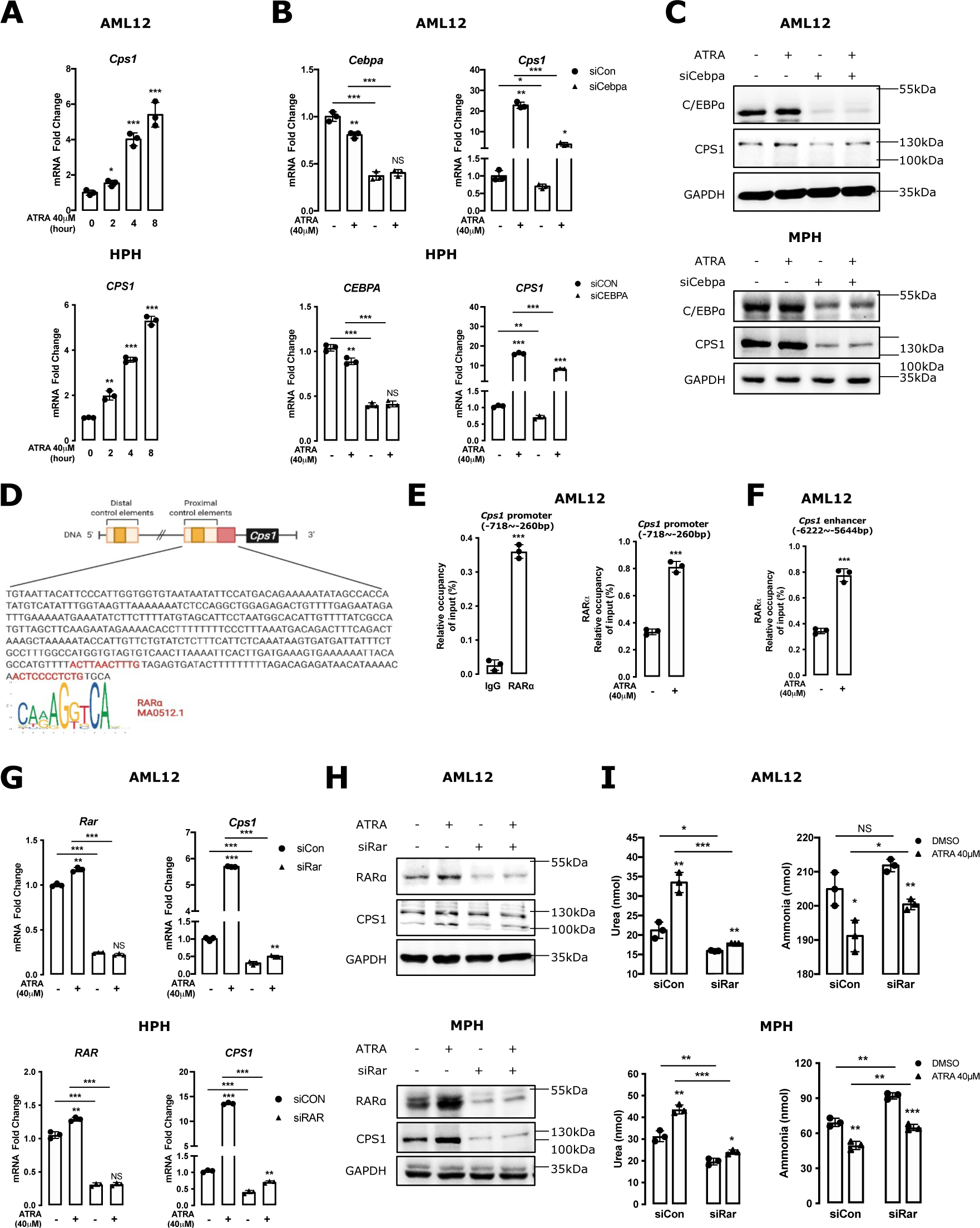
RAR is a key factor to regulate urea cycle in hepatocytes lacking C/EBPα. **(A)** qPCR analyzed the effects of all-trans-retinoic acid (ATRA) (40μM) on mRNA expression of CPS1 in AML12 cells and human primary hepatocytes (HPH) at different time points. **(B-C)** qPCR **(B)** and western blotting **(C)** analyzed the effects of ATRA on CPS1 expression in AML12 cells and HPH or mouse primary hepatocytes (MPH) after Cebpa siRNA transfection for 24h. **(D)** The binding sites of RARα on *Cps1* (-718∼-260bp) promoter were predicted by Jaspar dataset (*26*). The putative RARα binding sites are dyed red. **(E)** ChIP-qPCR analyzed the binding activities of RARα on *Cps1* (-718∼-260bp) promoter in AML12 cells (left panel) or treated with 40μM ATRA for 24 hours (right panel). **(F)** ChIP-qPCR also analyzed the effect of ATRA on RAR binding on the region of *Cps1* enhancer (-6222∼-5644bp). **(G-H)** After Rar siRNA transfection for 24h, AML12 cells were treated by ATRA for additional 24h. qPCR **(G)** and western blotting **(H)** were performed to analyze the expression of RARα and CPS1. **(I)** Urea and ammonia levels were examined in the culture medium of AML12 cells and MPH transfected with Rar siRNA for 24 hours, followed by ATRA treating for another 24 hours. Bars represent the mean ± SD, P values were calculated by One-way ANOVA, *, P<0.05; **, P<0.01, and ***, P<0.001 and NS, no significance. Triple experiments were performed, and one representative result is shown.

Besides the *Cps1* promoter, the binding of RAR to the *Cps1* enhancer was also examined. ChIP assays confirmed that RAR was capable of binding to the *Cps1* liver-specific element localized 6kbp upstream of the transcription start site, where FOXA2, H3K4me1 and H3K27ac binding were enriched (**Figure 3B, 5F**) and C/EBPα also bound (**Figure 3C**).

Knockdown of RAR completely inhibited ATRA-induced Cps1 expression in hepatocytes (**Figure 5G-H**). Consistent with increased Cps1 expression, ATRA increased urea levels and reduced ammonia concentrations in AML12 cells and MPHs (**Figure 5I**). The reduction in ammonia required RAR, given that knockdown of *Rar* significantly inhibited the effect (**Figure 5I**).

These results suggest a crucial role of RAR in rescuing urea cycle gene expression in hepatocytes that have lost C/EBPα expression.

### Foxa2-AAV8 improves urea cycle and reduces serum ammonia concentrations in acetaminophen-treated mice

Given that dysregulation of urea cycle and hyperammonemia occur in acetaminophen (APAP)-induced ALF, the ALF model through injecting APAP (300mg/kg body weight) into mice was established. The liver tissues at 3h, 6h, 12h and 24h following APAP injection were examined. H&E staining demonstrated rapidly increased liver injury and necrosis over time (**Figure S1A**). At 3h, the damaged livers showed necrotic areas surrounding central veins. At 24h, all examined livers showed massive hepatic necrosis (MHN) (**Figure S1A**). Consistent with liver damage, serum ALT and AST concentrations also remarkably increased with time (**Figure S1B**). Following APAP injection, serum urea levels were significantly reduced at any detected time point (**Figure S1C**). Peak serum ammonia levels occurred at 3h after APAP administration (**Figure S1C**). High serum ammonia levels continued at 6h following APAP insult and reduced to normal levels 12h later (**Figure S1C**).

To evaluate the effects of FOXA2 on the maintenance of urea cycle in ALF, Foxa2-AAV8 were injected into mice. qPCR, western blotting and IF analyses confirmed increased Foxa2 expression and location of Foxa2-AAV8 in hepatocytes (**Figure 6A-C**). Serum and liver tissues of mice at 3h following APAP administration were examined. Injection of Foxa2-AAV8 reduced APAP-induced serum ALT and AST concentrations (**Figure 6D**). Histologically, Foxa2-AAV8 reduced APAP-induced necrotic areas (**Figure 6E**). Impressively, administration of Foxa2-AAV8 maintained serum urea and ammonia levels at the normal levels in APAP-treated mice (**Figure 6F**). qPCR analyses showed that hepatic Cps1 mRNA expression was significantly increased in Foxa2-AAV8-treated mice (**Figure 6G**).

**Fig. 6.**
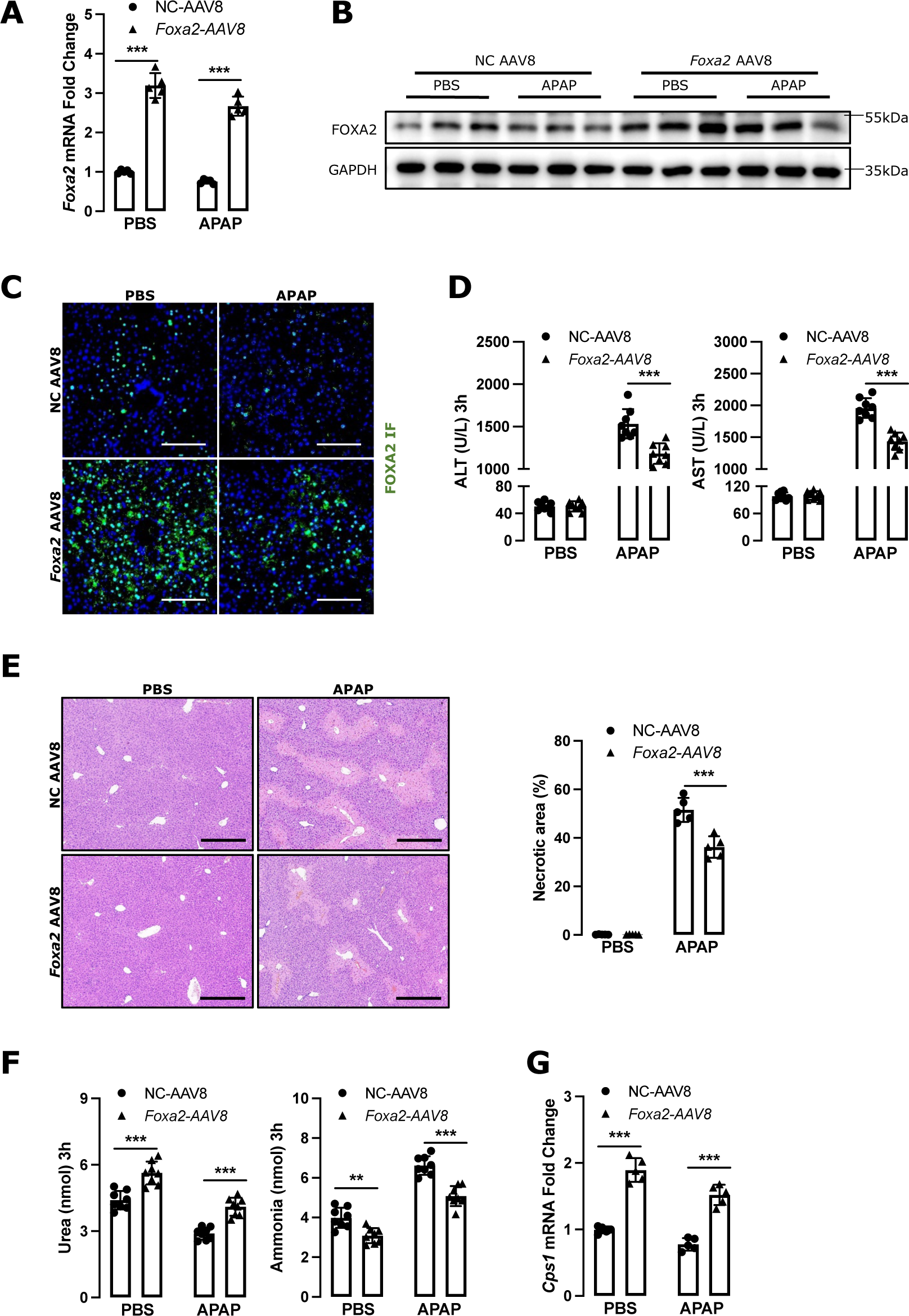
Foxa2-AAV8 improves urea cycle and prevents hyperammonemia in acetaminophen-treated mice. **(A)** qPCR, western blotting **(B)** and **(C)** immunofluorescence showed the mRNA, protein expression and location of Foxa2 in the PBS- or acetaminophen (APAP)-treated mice, which were injected with vector-AAV8 or Foxa2-AAV8. **(D)** ALT, AST, **(F)** urea and ammonia concentrations in serum samples were collected at 3 hours after APAP injection and were analyzed as described in Materials and Methods section. **(E)** H&E staining showed the necrotic areas in the live tissues of mice with indicated administration. Necrotic area was quantified. **(G)** qPCR determined the mRNA expression of Cps1 in the mouse liver tissues. Bars represent the mean ± SD, P values were calculated by One-way ANOVA, **, P<0.01, and ***, P<0.001. Triple experiments were performed, and one representative result is shown.

These results suggest a crucial role of FOXA2 in the maintenance of urea cycle and preventing hyperammonemia in ALF.

### C/EBPα and FOXA2 regulate *CPS1* transcription and urea cycle in liver progenitor cells

MHN is the most severe histological alteration in ALF (*9*). Interestingly, not all ALF patients with MHN suffer from HE. How can these ALF patients perform urea cycle to prevent HE in the absence of sufficient hepatocytes? LPCs might be the leading player that takes over urea cycle to maintain ammonia homeostasis in these patients.

Next, the expression of urea cycle relevant genes was compared between hepatocytes and LPCs. RNA-seq analyses showed that all six urea cycle enzyme transcripts were higher in human and mouse hepatocytes than in LPC lines, BMOL and HepaRG cells (**Figure 7A**). As in hepatocytes, FOXA2 bound to the *Cps1* promoter in BMOL cells (**Figure 7B**). In contrast to hepatocytes, LPCs usually do not express C/EBPα (*6*). In hepatocytes, CPS1 was constitutively expressed given that Pol II with phosphorylation on serine 5 (S5) and serine 2 (S2) of the heptapeptide repeats in the C-terminal domain of the Rbp1 subunit bound to the *Cps1* core promoter (**Figure S2A**). However, both Pol II S5 and Pol II S2 did not bind to the promoter of *Cps1* gene in LPCs (**Figure S2B**). Therefore, normal LPCs do not express CPS1. Impressively, overexpression of *Foxa2* increased the binding of Pol II S5 and Pol II S2 to the *Cps1* core promoter (**Figure 7C**). Furthermore, *Foxa2* overexpression increased urea levels and reduced ammonia concentrations in supernatant of LPCs, whereas knockdown of *Foxa2* reduced urea concentrations, but increased ammonia levels (**Figure 7D**). As in hepatocytes, qPCR analysis revealed that overexpression of Foxa2 did not significantly increase mRNA expression of Cps1 in LPCs (**Figure 7E**), indicating a pioneer factor-like effect.

**Fig. 7.**
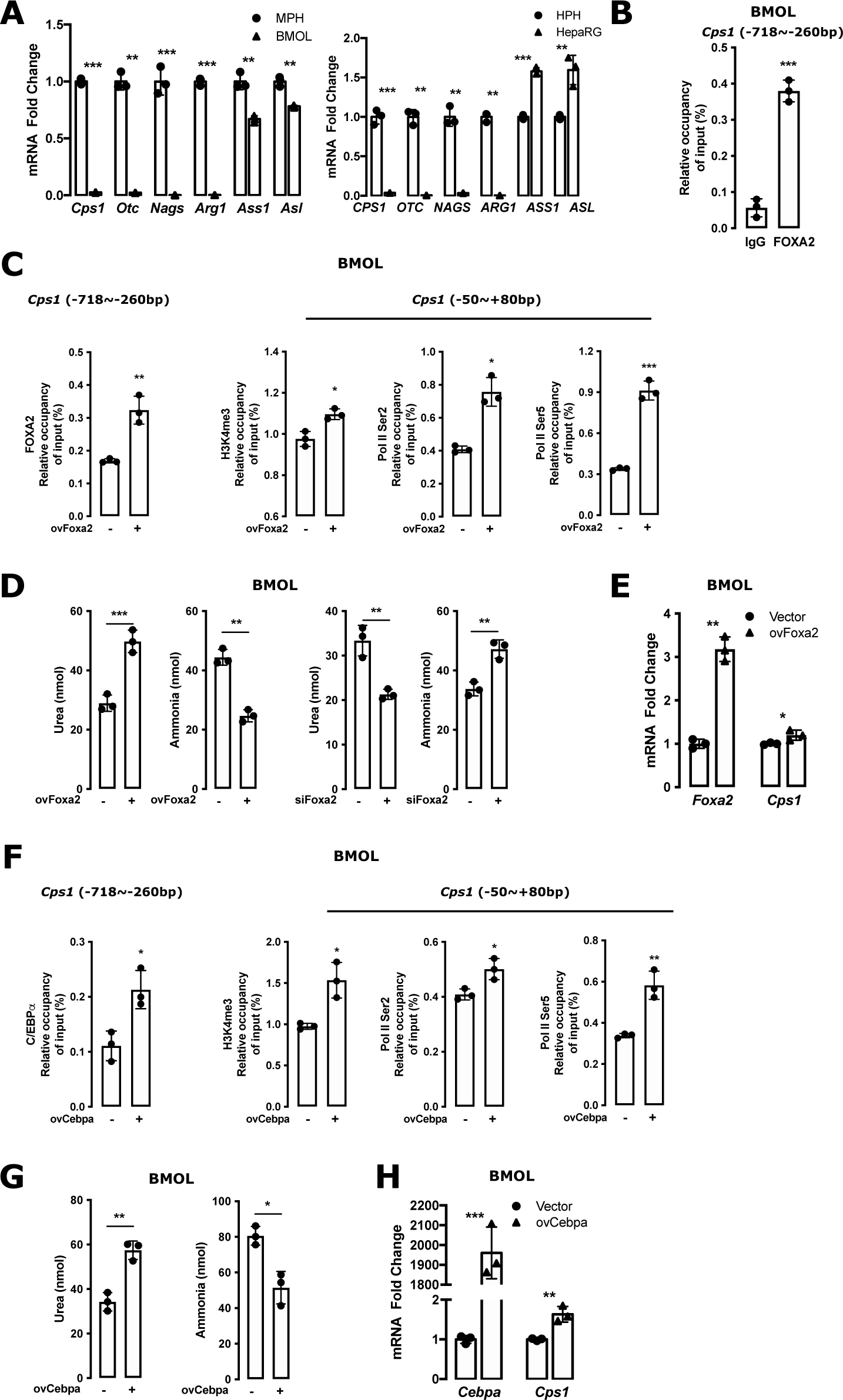
FOXA2 and C/EBPα regulate CPS1 transcription and urea cycle in liver progenitor cells. **(A)** mRNA expression of urea cycle enzymes in liver progenitor cells and hepatocytes (MPH vs. BMOL, HPH vs. HepaRG) was analyzed by qPCR. **(B)** ChIP-qPCR examined FOXA2 binding to the *Cps1* (-718∼-260bp) promoter in BMOL cells. **(C)** ChIP-qPCR analyzed the binding activities of FOXA2 on *Cps1* promoter (-718∼-260bp), and H3K4me3, Pol II S2 and Pol II S5 on *Cps1* core promoter (-50∼+80bp) in BMOL cells treated with Foxa2 construct for 48 hours. **(D)** Urea and ammonia levels were measured in the culture medium of BMOL cells transfected with Foxa2 construct or siRNA for 48 hours. **(E)** qPCR analyzed the effects of Foxa2 on Cps1 expression in BMOL cells with Foxa2 constructs. **(F)** ChIP-qPCR examined the binding activities of C/EBPα on *Cps1* promoter (-718∼-260bp), and H3K4me3, Pol II S2 and Pol II S5 on *Cps1* core promoter (-50∼+80bp) in BMOL cells treated with Cebpa construct for 48 hours. **(G)** Urea and ammonia levels were examined in the culture medium of BMOL cells with Cebpa construct for 48 hours. **(H)** qPCR analyzed the effects of C/EBPα on Cps1 expression in BMOL cells with Cebpa construct for 48 hours. Bars represent the mean ± SD, P values were calculated by unpaired Student’s t-test, *, P<0.05; **, P<0.01, and ***, P<0.001 and NS, no significance. Triple experiments were performed, and one representative result is shown.

Although LPCs do not express C/EBPα physiologically, robust C/EBPα expression in LPCs can be detected in cirrhotic patients with ductular reaction (**Figure S3**). These LPCs also expressed CPS1 (**Figure S3**), indicating a potential association between C/EBPα expression and urea cycle enzyme synthesis in LPCs in vivo. To confirm this association, we forced transgenic ectopic Cebpa expression in LPCs in vitro. ChIP assays showed that overexpression of *Cebpa* increased the binding of H3K4me3, Pol II S5 and S2 to the *Cps1* core promoter (**Figure 7F**). Overexpression of *Cebpa* increased Cps1 mRNA expression and urea levels, but reduced ammonia concentrations in supernatant of LPCs (**Figure 7G-H**).

These results suggest that as an alternative primary cell, LPCs are capable of exploiting a similar transcription network to perform urea cycle in the absence of hepatocytes.

## DISCUSSION

To date, the transcriptional regulation of urea cycle genes to prevent the occurrence of HE in different pathophysiological conditions, including ALF, have been largely unknown. The current study provides some molecular details of the CPS1 transcriptional regulatory mechanisms in physiological condition and ALF. In normal hepatocytes, RNA Pol II binds to the *Cps1* core promoter, which results in constitutively expression of CPS1. In addition, sufficient CPS1 transcription requires contribution from FOXA2 and C/EBPα. FOXA2 does not only bind directly to the *CPS1* promoter, but also acts as a pioneer factor to maintain chromatin accessibility of the gene enhancers, providing open binding sites for additional transcription regulators such as C/EBPα. In ALF, hepatic C/EBPα expression is inhibited by inflammatory cytokines such as TGF-β and TNF-α (*10*). In this setting, nuclear receptor RAR replaces C/EBPα and synergizes with FOXA2 to maintain CPS1 transcription in hepatocytes. Once ALF patients suffer substantial loss of functional hepatocytes through MHN, LPCs uphold the vital urea cycle function of metabolizing ammonia through initiating the transcription network described above, comprising FOXA2 and C/EBPα. This compensatory mechanism rescues ALF patients from HE. It is remarkable that the same “rescue network” involving FOXA2 and C/EBPα is also in charge of ALB transcription in ALF (*6*). Whether this regulatory network is a general hub in charge of vital hepatic functional genes under pathological conditions requires further investigation.

As pioneer factor, all FOXA family members FOXA1, FOXA2 and FOXA3 play a crucial roles in liver development and in maintaining essential liver functions in the adult liver (*11*). Ablation of all FOXA genes in adult liver results in collapse of essential hepatic gene regulatory networks and kills mice (*12*). In the absence of FOXA genes, the urea cycle in mouse liver is disrupted, resulting in low urea levels and hyperammonemia (*12*). In ALF patients described in a previous study, only FOXA2 expression is detectable by immunohistochemistry in a portion of liver tissues (*13*). This study further shows that robust FOXA2 is expressed in the ALF patients without HE, but not in those with HE. More impressively, FOXA2 expression is associated with CPS1 levels in both hepatocytes and LPCs, indicating that FOXA2 might regulate CPS1 transcription in ALF. FOXA2 possesses binding sites on the *CPS1* enhancer 6.3kb upstream of the transcription starting site (*4*). This region also contains C/EBPα binding sites (*4*). The current study clarifies the role of FOXA2 and C/EBPα in the regulation of *CPS1* transcription: FOXA2 acts as a pioneer factor to maintain chromosome accessibility of the *CPS1* enhancer, which is required for C/EBPα binding to the enhancer.

In ALF, severe inflammation profoundly influences the transcriptional regulatory network that maintains urea cycle. Previous studies showed that FOXA2 expression is upregulated by multiple factors such as TNF-α, hedgehog and glucagon (*6*, *13*). Therefore, hepatic FOXA2 expression is remarkably induced in the condition of inflammatory response. In contrast to FOXA2, C/EBPα is rapidly inhibited by inflammatory cytokines such as TGF-β and TNF-α (*6*). How is the urea cycle maintained in the absence of C/EBPα? We observed that RAR is the “deputy transcription factor” that maintains CPS1 transcription through binding to their enhancers. This type of compensatory rescue mechanism has previously been shown in the ALB transcription (*6*).

Given the key roles of FOXA2 and C/EBPα in the maintenance of urea cycle, could pharmaceutical induction of these factors might represent a potential approach to treat HE in ALF? Animal experiment based on APAP-induced mouse ALF confirmed that injection of Foxa2-AAV8 maintains urea cycle and prevents the occurrence of hyperammonemia. In a recent study, ectopic Foxa2 expression by adenovirus restores apical multidrug resistance protein 2 expression and normalized serum bilirubin levels in lipopolysaccharide-treated mice and farnesoid X receptor knockout mice (*13*). In addition, FOXA2 expression is essential for ALB expression in ALF (*6*). These results suggest that pharmaceutical induction of FOXA2 might be a promising new approach to treat ALF. One major concern is whether ectopic expression of a pioneer factor such as FOXA2 might as well open chromatin in the enhancers of unwanted genes, with a plethora of unintended consequences. However, in an emergency such as HE in ALF, the risk-benefit balance might justify a temporary intervention that would be unacceptable in a more chronic treatment regime.

In addition to FOXA2, this study also investigated the effects of ectopic Cebpa expression on APAP-induced ALF in mice. Injection of Cebpa-AAV8 significantly induced severe inflammation and increased liver injury in mice (data not shown). This effect might be due to the effects of C/EBPα on inflammatory cells, where C/EBPα also acts as a pioneer factor: C/EBPα reprograms B cell differentiation into macrophages (*14*). When Cebpa-AAV8 is injected into mice, besides entering hepatocytes, Cebpa-AAV8 also localizes to inflammatory cells and induces severe inflammatory response in APAP-treated mice (data not shown). Further experiments to determine the usefulness of pharmaceutical induction of hepatic C/EBPα in treating HE in ALF are expected in the near future.

Apart from stable transcriptional regulation in hepatocytes, the maintenance of an active urea cycle in LPCs is crucial for the survival of ALF patients who suffer from MHN. This study found that LPCs are capable of initiating transcription of a network comprising FOXA2 and C/EBPα to ensure ammonia metabolism in ALF. Physiologically, LPCs do not express detectable FOXA2 and C/EBPα (*6*). In ALF patients, particularly in those with good prognosis, both factors are robustly expressed in both LPCs and remaining hepatocytes (*6*). How are FOXA2 and C/EBPα activated in LPCs? Previous studies showed that expression of FOXA2 in LPC is induced by factors such as TNF-α, glucagon and hedgehog (*6*, *13*). To date, it is unknown if the same factors govern the expression of C/EBPα in LPCs. Given the lack of an ideal animal model to scrutinize LPC activation and functions in ALF, clarification of these issues is a challenge in understanding the detailed molecular mechanisms of ALF. Based on current knowledge, a hierarchical transcriptional network that regulates urea cycle is highlighted in **Figure 8**.

**Fig. 8.**
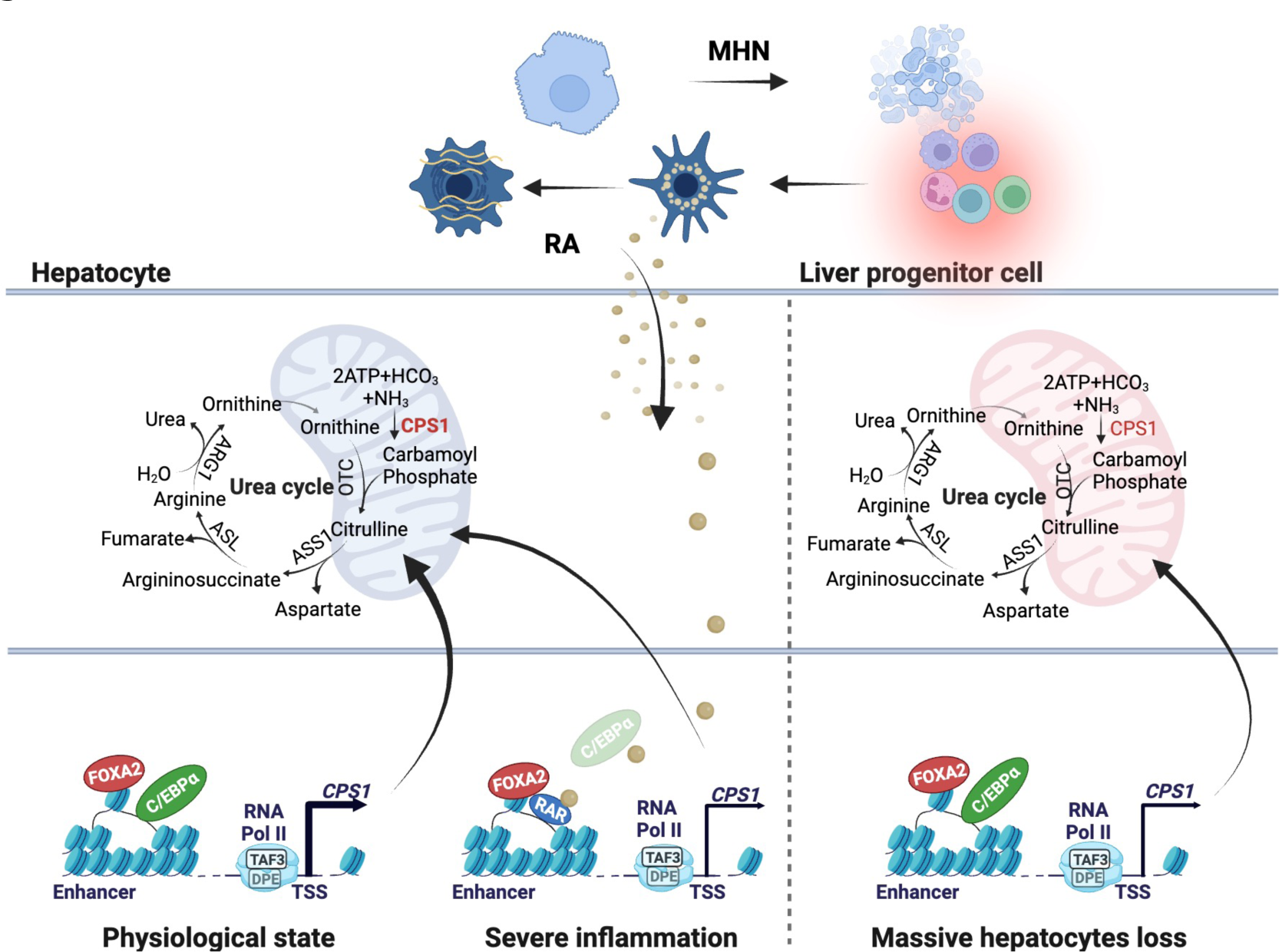
Schematic diagram shows a hierarchical network that maintains urea cycle in ALF. (1) Physiologically, CPS1 transcription requires pioneer factor FOXA2 to maintain chromatin accessibility on its enhancer, which is essential for C/EBPα binding to DNA and activate gene transcription. (2) When hepatic C/EBPα expression is inhibited by inflammation, retinoic acid receptor (RAR) synergizes with FOXA2 to maintain CPS1 transcription. (3) Once ALF patients suffer from massive hepatic necrosis, liver progenitor cells (LPCs) perform the urea cycle to prevent hyperammonemia by initiating a transcription network comprising FOXA2 and C/EBPα.

## MATERIALS AND METHODS

### Study design

A total of 35 patients with ALF were enrolled in the Department of Medicine II, University Medical Center Mannheim, Medical Faculty Mannheim, Heidelberg University and the Department of Gastroenterology and Hepatology, Beijing You’an Hospital, Affiliated with Capital Medical University (*6*). Among these patients, 15 were diagnosed as HE according to West-Haven criteria (*15*). Liver tissues were collected when patients received tumor excision or liver transplantation. ALF is defined as a severe liver injury, leading to coagulation abnormality usually with an international normalized ratio ≥ 1.5, and any degree of mental alteration (encephalopathy) in a patient without pre-existing liver disease or with an illness of up to 4 weeks duration (*1*). The study protocol was approved by local ethics committees (Jing-2015-084, and 2017-584N-MA). Written informed consent was obtained from patients or their representatives. Detailed patient information is given in **Table S2**.

### Cells

AML12 cells were grown in DMEM/F-12 medium (21331-020, Gibco) supplemented with 10% fetal bovine serum (FBS), 5% Insulin-Transferrin-Selenium, 2 mM L-glutamine, penicillin (100 U/mL)-streptomycin (100 μg/mL) (P/S), and 100 nM dexamethasone. Mouse primary hepatocytes (MPHs) were isolated from ad libitum fed mice using the two-step collagenase perfusion technique as described previously (*16*),(*17*). The isolated cells were seeded on collagen-coated plates in Williams’ E medium supplemented with 10% FBS, 2mM L-glutamine, 1% P/S, 100 nM dexamethasone, and 5% Insulin-Transferrin-Selenium. Human primary hepatocytes (HPHs) were isolated by the Cell Isolation Core Facility of the Biobank Großhadern, University Hospital, LMU Munich. BMOL cells were maintained in Williams’ E medium supplemented with 10% FBS, 2mM L-glutamine, 1% P/S, and 10 μg/mL Insulin (I0516, Sigma-Aldrich). HepaRG cells were purchased from Biopredic International (Saint Gregoire, France). The cells were grown in Williams’ E medium supplemented with 10% FBS, 2mM L-glutamine, 1% P/S, 5 μg/ml insulin, and 50 μM hydrocortisone hemisuccinate (PHR1926, Sigma-Aldrich). All cells were cultured at 37°C in a humidified atmosphere containing 5% CO2.

### Animal experiment

C57BL/6J mice were purchased from the Gempharmatech Co., Ltd, China. Animals were housed in temperature- and light/dark cycle-controlled rooms. Mice received 2.0x1011 viral genomes of adeno-associated virus serotype 8 (AAV8)-Foxa2-GFP (n=16) or AAV8-GFP (n=16) through the tail vein. After 14 days, half of the mice were administered intraperitoneally with acetaminophen (APAP) at a dose of 300 mg/kg after overnight fasting (16 hours), and then resumed feeding. The samples were collected at different timepoints. All the animal experimentation was conducted in accordance with the guidelines of the animal ethics committee of Anhui Medical University.

### Immunohistochemistry

Liver tissue sections were dewaxed in xylene (9713.2, Carl Roth) and rehydrated in an ethanol gradient (K928.4, Carl Roth). For antigen retrieval, slides incubated with 1 mM Ethylene Diamine Tetraacetic Acid solution (EDTA, E9884, Sigma-Aldrich, pH 8.4) or citrate acid buffer (C2404, Sigma-Aldrich, pH 6.0) were heated by microwave (1000 W, 15 seconds full power, 45 seconds shut down, 10 cycles). Following cooling at room temperature for 30 mins, the slides were incubated in DAKO blocking peroxide (S200389-2, DAKO) for 30 mins to eliminate non-specific staining. Subsequently, primary antibodies (**Table S3**) were incubated overnight at 4°C. Next day, following 3 washes with phosphate-buffered saline (PBS, L182-50, Biochrom), sections were incubated with the horseradish peroxidase-labeled secondary antibodies (**Table S3**) for 45 mins. Visualization was achieved with 3,3-diaminobenzidine (DAB, D5905, Sigma-Aldrich). Counterstaining was performed with hematoxylin (H-3401-500, Vector Laboratories). Finally, all sections were dehydrated and mounted with mounting medium (C9368, Sigma-Aldrich). Images were taken under a microscope (DMRBE, Leica).

### H&E staining

Four-micrometer-thick sections from paraffin-embedded liver tissues were fixed in 4% paraformaldehyde. The slides were deparaffinized and rehydrated with distilled water. After staining in hematoxylin for 1 min, the sections were washed in tap water for 10 mins. After counterstaining in alcohol-eosin for 1 min, the slides were dehydrated through 3 changes of 95% EtOH, 2 changes of 100% EtOH and 3 changes of Xylene for 1 min each. Images of mounted sections were taken under a microscope.

### Chromatin immunoprecipitation and quantitative real-time PCR

Chromatin immunoprecipitation and quantitative real-time PCR (ChIP-qPCR) analyses were performed as described previously, with minor modifications (*18*). Briefly, for each ChIP, cells were isolated from one 10 cm cell culture dish (95% confluence). Cells were washed twice with PBS (14190144, Thermo Fisher) and incubated with 1% formaldehyde at 37 °C for 10 mins to crosslink proteins and DNA. Following ice-cold PBS washing twice, the cells were collected in lysis buffer (1% SDS, 5 mM EDTA in 50 mM Tris, pH 8.1) containing 1% protease inhibitor cocktail. The chromatin was sonicated by 30-seconds pulses, followed by 30-seconds interval to produce DNA fragments with an average of 200-500 bp length. After sonication, the samples were centrifuged at 14000 rpm, 4°C for 10 mins. The supernatants were collected in a clean tube and diluted by adding dilution buffer. 5μg of the antibody or normal IgG and 50μL of agarose beads were added into the samples. After rotating overnight at 4°C, the beads were collected by centrifugation at 2000 rpm for 10 mins. Then the pellets were successively washed with 1 ml TSEI (0.1% SDS, 1% Triton X-100, 2 mM EDTA, 20 mM Tris-HCl Ph8.0, 150 mM NaCl), TSEII (0.1% SDS, 1% Triton X-100, 2 mM EDTA, 20 mM Tris-HCl Ph8.0, 500 mM NaCl), BufferIII (0.25 M LiCl, 1% NP-40, 1% Deoxycholic acid, 10 mM Tris-HCl Ph8.0) and TE buffer (10 mM Tris-HCl Ph8.0, 1 mM EDTA) for 10 minutes each at 4°C. After centrifugation at 2000 rpm for 10 minutes, the beads were resuspended in elution buffer and kept washing at room temperature for 10 mins. Immunoprecipitated complexes were de-crosslinked by heating to 65 °C for 6h. The pulled-down DNA fragments were extracted and purified using phenol-chloroform extraction/ethanol precipitation. The samples were subjected to quantitative PCR using SYBR green assay (A25918, Thermo Fisher). The primer sets used for these assays are listed in **Table S4**.

### RNA extraction and quantitative real-time PCR

All reagents and containers used for RNA processing were RNase-free grade or treated with 0.1% DEPC (4387937, Thermo Fisher) to eliminate RNase contaminants. Total RNA was extracted from cells and tissues in Trizol (15596018, Invitrogen) according to the manufacturer’s instructions. Reverse transcription of 500 ng total RNA to synthesize cDNA was performed using random primers (SO142, Thermo Fisher) and RevertAid H Minus Reverse Transcriptase (EP0452, Thermo Fisher) according to the manufacturer’s instructions. The qPCR assays were carried out on a StepOnePlus system (Applied Biosystem) using SYBR Green Master Kit (A25918, Thermo Fisher). Primers for qPCR are listed in **Table S4**. Three biological replicates per condition were measured. The relative abundance (fold changes) of each target gene compared with a set of internal controls were determined by the -2ΔΔCT method (*19*).

### Western Blotting

Cells were lysed in RIPA buffer (R0278, Sigma-Aldrich) supplemented with protease inhibitors (36978, Thermo Fisher) and phosphatase inhibitors (P5726, Sigma-Aldrich) on ice for 10 mins. Cell lysates were collected by centrifugation at 14000 rpm for 15 mins at 4°C. Total protein concentrations in the supernatant were determined by a protein assay kit (5000113-115, Bio Rad, Bradford method). Following the addition of LDS sample buffer (4x, NP0007, Lifetechnologies), the samples were boiled at 90°C for 10 mins. Twenty microgram proteins were separated by 10-15% SDS-PAGE and transferred onto 0.2 μm nitrocellulose membranes (4685.1, Carl Roth). Following blocking with 5% non-fat milk for 1 hour at room temperature, the membranes were incubated with primary antibodies (**Table S3**) at 4°C overnight. Next day, the membranes were washed with tris buffered saline plus 0.05% Tween 20 (TBST) and incubated with secondary antibodies (**Table S3**) for 1 hour. After washing with TBST three times, the membranes were incubated and visualized by the western blotting substrate (32132, Lifetechnologies).

### siRNA transfection

Confluent cells (80% primary hepatocytes or 60% cell lines) were seeded 24 hours before transfection. Small interfering RNAs (siRNAs) (**Table S5**) were transfected into cells using Lipofectamine RNAiMAX reagent (13778-075, Thermo Fisher) according to the manufacturer’s instruction. The transfected cells were incubated at 37 °C for 48 hours, followed by extraction of cellular DNA, RNA, and proteins. A non-targeting negative stealth siRNA (scrambled, sc-37007, Santa Cruz) was used as a negative control. The sequences of siRNA are listed in **Table S6**.

### Transient transfection assay

Transfection was performed using lipofectamine 3000 reagent (L3000008, Thermo Fisher) according to the manufacturer’s instructions. Briefly, cells were seeded in 12-well plates and grown to 90∼95% confluency. 1.75 μg/well of plasmids (**Table S5**) were transfected into cells. After 48 hours, the cells were washed with PBS. Cellular supernatant, DNA, RNA, and proteins were collected.

### Measurement of urea and ammonia concentrations

The urea and ammonia assay kits were purchased from Abcam (Ab83362, Ab83360). 2.5 x 10^5^ cells were resuspended in 2 mL complete medium and added to a 6-well plate. Cells were incubated at 37 °C overnight, then the supernatant was replaced with the fresh medium, followed by the corresponding treatment. 10 μl supernatant or serum was diluted to 50 μl volume with the assay buffer and used for the detection assay. 50 μl of reaction mix was added into each standard and sample wells. The plate was incubated at 37 °C for 60 mins protected from light. Urea production and ammonia levels in the medium were determined by measuring absorbance at 570 nm. Meanwhile, the standard curve was produced using a series of urea or NH4Cl standards with different concentrations, which were used to calculate the urea or ammonia concentration in the samples.

### Serum ALT and AST measurement

The levels of serum ALT and AST were measured by Mairui automated biochemical analyzer (BS-350e, China). The values were expressed as U/L of sera.

### RNA sequencing

Primary hepatocytes were freshly isolated from three mice. MPHs were transfected by Cebpa siRNA for 48 hours. HPHs, MPHs and LPCs (HepaRG and BMOL cells) were stably cultured in the indicated culture medium (described above) before RNA collection. Total RNA was extracted as described above. RNA quality was checked with the Agilent 2100 Bioanalyzer and the RNA 6000 Nano Kit (Agilent, Waldbronn). Samples with RNA integrity number above 9.5 were used for RNA sequencing. The sequencing work was performed by BGI Tech Solutions Co. (Hong Kong, China).

### Bioinformatic analyses of RNA sequencing

Most of the analysis was performed with R and bioconductor using the NGS analysis package systempipeR (*20*). Quality control of raw sequencing reads was performed using FastQC (Babraham Bioinformatics). Low-quality reads were removed using trim-galore (version 0.6.4). The resulting reads were aligned to human genome version GRCh38.p12 and mouse genome version GRCm38.p6 from Gene code and counted using Kallisto (version 0.46.1) (*21*). The count data was transformed to log2-counts per million using the voom-function from the limma package (*22*). Differential expression analysis was performed using the limma package in R. A false positive rate of α= 0.05 with FDR correction was taken as the level of significance. Volcano plots and heatmaps were created using the ggplot2 package (version 2.2.1) and the complex heatmap (version 2.0.0) (*23*).

### Transcript Profiling

RNA sequencing data used in this study were obtained from the Gene Expression Omnibus (GEO) repository (accession no. GSE 181201) (*6*).

### Data Availability

ChIP sequencing (ChIP-seq) data of mice liver tissue were obtained from the GEO (accession no. GSE 157452, GSE65167) (*24*, *25*).

### Statistical analysis

Analyses were performed using SPSS Statistics 23.0. The unpaired Student’s t-test was used to determine statistically significant differences between two groups. P values less than 0.05 were considered significant and represented graphically as *, P<0.05; **, P<0.01; and ***, P<0.001.

## List of Supplementary Materials

Table S1. Immunohistochemical staining for CPS1, FOXA2 and C/EBPα in 20 ALF patients*

Table S2. Characteristics of the study population

Table S3. Antibodies used in the study

Table S4. Primers used in the study

Table S5. siRNA and plasmid used in the study

Table S6. The sequences of siRNA

Table S7. Reagent kits used in the study

Fig. S1. Acetaminophen injections induce liver injury in mice at different time points

Fig. S2. The analysis of Cps1 core promoters in hepatocytes and liver progenitor cells

Fig. S3. C/EBPα and CPS1 expression in cirrhotic patients with ductular reactions

References (*1–2*)

## Acknowledgments

We are grateful to Drs. Nelson Fausto and George Yeoh for kindly providing AML12 and BMOL cells. We thank the Human Tissue and Cell Research Foundation, a nonprofit foundation regulated by German civil law, which facilitates research with human tissue through the provision of an ethical and legal framework for prospective sample collection. We acknowledge the support of the LIMA Live Cell Imaging at Microscopy Core Facility Platform Mannheim (CFPM).

## Funding

Deutsche Forschungsgemeinschaft WE 5009/9-1, 5009/12-1, DO 373/24-1 (H.L.W.)

Chinese-German Cooperation Group projects GZ 1517 (H.L.W., H.D.), M-0099 (S.D., H.W.), M-0200 (S.S.W., C.M.)

Chinese Nature Science Foundation 81970525 (H.D.), 81870424 (S.S.W.), 82300699 (R.L.F)

Beijing Natural Science Foundation Program and Scientific Research Key Program of Beijing Municipal Commission of Education KZ201810025037 (H.D.)

LiSyM Grant PTJ-FKZ: 031L0043 (S.D.)

Beijing Municipal Natural Science Foundation 7212052 (S.S.W.) BMBF through HiChol 01GM1904A (R.L.)

Shanghai Rising-Star Program 23YF1423200 (R.L.F)

Chinese Scholarship Council 202106320043 (C.T.)

## Author contributions

Conceptualization: H.L.W.

Methodology: R.L.F., Ru.L., C.H.T., Y.J.L., S.M., S.W., X.F.L., U.W., H.N.,

Investigation: R.L.F., Ru.L., H.L., C.S.,

Visualization: R.L.F., Ru.L., T.L., K.J.K., Ca.S.

Funding acquisition: H.L.W., H.G.D., S.D., S.S.W., R.L.F., Ro.L., C.H.T., C.M.

Project administration: H.L.W., H.W.

Supervision: H.L.W., H.W.

Writing – original draft: R.L.F., H.L.W.

Writing – review & editing: R.L.F., C.M., Ro.L., M.P.A., S.D., H.G.D., H.W., H.L.W.

## Data and materials availability

All data are available in the main text or the supplementary materials.

## References and Notes

1. J. G. O’Grady, S. W. Schalm, R. Williams, Acute liver failure: redefining the syndromes. Lancet. 342, 273–275 (1993).

2. J. Vaquero, C. Chung, M. E. Cahill, A. T. Blei, Pathogenesis of hepatic encephalopathy in acute liver failure. Semin Liver Dis. 23, 259–269 (2003).

3. V. Walker, Ammonia metabolism and hyperammonemic disorders. Adv Clin Chem. 67, 73– 150 (2014).

4. Y.-R. Chen, K. Sekine, K. Nakamura, H. Yanai, M. Tanaka, A. Miyajima, Y-box binding protein-1 down-regulates expression of carbamoyl phosphate synthetase-I by suppressing CCAAT enhancer-binding protein-alpha function in mice. Gastroenterology. 137, 330–340 (2009).

5. T. Kimura, V. M. Christoffels, S. Chowdhury, K. Iwase, H. Matsuzaki, M. Mori, W. H. Lamers, G. J. Darlington, M. Takiguchi, Hypoglycemia-associated hyperammonemia caused by impaired expression of ornithine cycle enzyme genes in C/EBPalpha knockout mice. J Biol Chem. 273, 27505–27510 (1998).

6. R. Feng, K. Kan, C. Sticht, Y. Li, S. Wang, H. Liu, C. Shao, S. Munker, H. Niess, S. Wang, C. Meyer, R. Liebe, M. P. Ebert, S. Dooley, H. Ding, H. Weng, A hierarchical regulatory network ensures stable albumin transcription under various pathophysiological conditions. Hepatology (2022), doi:10.1002/hep.32414.

7. T. Lin, R. Feng, R. Liebe, H.-L. Weng, Liver Progenitor Cells in Massive Hepatic Necrosis-How Can a Patient Survive Acute Liver Failure? Biomolecules. 12, 66 (2022).

8. M. Iwafuchi-Doi, G. Donahue, A. Kakumanu, J. A. Watts, S. Mahony, B. F. Pugh, D. Lee, K. H. Kaestner, K. S. Zaret, The Pioneer Transcription Factor FoxA Maintains an Accessible Nucleosome Configuration at Enhancers for Tissue-Specific Gene Activation. Mol Cell. 62, 79– 91 (2016).

9. H.-L. Weng, X. Cai, X. Yuan, R. Liebe, S. Dooley, H. Li, T.-L. Wang, Two sides of one coin: massive hepatic necrosis and progenitor cell-mediated regeneration in acute liver failure. Front Physiol. 6, 178 (2015).

10. R. Feng, C. Tong, T. Lin, H. Liu, C. Shao, Y. Li, C. Sticht, K. Kan, X. Li, R. Liu, S. Wang, S. Wang, S. Munker, H. Niess, C. Meyer, R. Liebe, M. P. Ebert, S. Dooley, H. Wang, H. Ding, H.-L. Weng, Insulin Determines Transforming Growth Factor β Effects on Hepatocyte Nuclear Factor 4α Transcription in Hepatocytes. Am J Pathol, S0002-9440(23)00365-6 (2023).

11. K. S. Zaret, S. E. Mango, Pioneer transcription factors, chromatin dynamics, and cell fate control. Curr Opin Genet Dev. 37, 76–81 (2016).

12. Y. Reizel, A. Morgan, L. Gao, Y. Lan, E. Manduchi, E. L. Waite, A. W. Wang, A. Wells, K. H. Kaestner, Collapse of the hepatic gene regulatory network in the absence of FoxA factors. Genes Dev. 34, 1039–1050 (2020).

13. S. Wang, R. Feng, S. S. Wang, H. Liu, C. Shao, Y. Li, F. Link, S. Munker, R. Liebe, C. Meyer, E. Burgermeister, M. Ebert, S. Dooley, H. Ding, H. Weng, Gut, in press, doi:10.1136/gutjnl-2022-326987.

14. C. van Oevelen, S. Collombet, G. Vicent, M. Hoogenkamp, C. Lepoivre, A. Badeaux, L. Bussmann, J. L. Sardina, D. Thieffry, M. Beato, Y. Shi, C. Bonifer, T. Graf, C/EBPα Activates Pre-existing and De Novo Macrophage Enhancers during Induced Pre-B Cell Transdifferentiation and Myelopoiesis. Stem Cell Reports. 5, 232–247 (2015).

15. H. Vilstrup, P. Amodio, J. Bajaj, J. Cordoba, P. Ferenci, K. D. Mullen, K. Weissenborn, P. Wong, Hepatic encephalopathy in chronic liver disease: 2014 Practice Guideline by the American Association for the Study of Liver Diseases and the European Association for the Study of the Liver. Hepatology. 60, 715–735 (2014).

16. P. O. Seglen, Preparation of isolated rat liver cells. Methods Cell Biol. 13, 29–83 (1976).

17. M. N. Berry, A. R. Grivell, M. B. Grivell, J. W. Phillips, Isolated hepatocytes--past, present and future. Cell Biol. Toxicol. 13, 223–233 (1997).

18. J. D. Nelson, O. Denisenko, K. Bomsztyk, Protocol for the fast chromatin immunoprecipitation (ChIP) method. Nat Protoc. 1, 179–185 (2006).

19. K. J. Livak, T. D. Schmittgen, Analysis of relative gene expression data using real-time quantitative PCR and the 2(-Delta Delta C(T)) Method. Methods. 25, 402–408 (2001).

20. T. W. H Backman, T. Girke, systemPipeR: NGS workflow and report generation environment. BMC Bioinformatics. 17, 388 (2016).

21. N. L. Bray, H. Pimentel, P. Melsted, L. Pachter, Near-optimal probabilistic RNA-seq quantification. Nat Biotechnol. 34, 525–527 (2016).

22. M. E. Ritchie, B. Phipson, D. Wu, Y. Hu, C. W. Law, W. Shi, G. K. Smyth, limma powers differential expression analyses for RNA-sequencing and microarray studies. Nucleic Acids Res. 43, e47 (2015).

23. Z. Gu, R. Eils, M. Schlesner, Complex heatmaps reveal patterns and correlations in multidimensional genomic data. Bioinformatics. 32, 2847–2849 (2016).

24. M. Qu, H. Qu, Z. Jia, S. A. Kay, HNF4A defines tissue-specific circadian rhythms by beaconing BMAL1::CLOCK chromatin binding and shaping the rhythmic chromatin landscape. Nat Commun. 12, 6350 (2021).

25. X. Sun, J.-C. Chuang, M. Kanchwala, L. Wu, C. Celen, L. Li, H. Liang, S. Zhang, T. Maples, L. H. Nguyen, S. C. Wang, R. A. J. Signer, M. Sorouri, I. Nassour, X. Liu, J. Xu, M. Wu, Y. Zhao, Y.-C. Kuo, Z. Wang, C. Xing, H. Zhu, Suppression of the SWI/SNF Component Arid1a Promotes Mammalian Regeneration. Cell Stem Cell. 18, 456–466 (2016).

26. J. A. Castro-Mondragon, R. Riudavets-Puig, I. Rauluseviciute, R. Berhanu Lemma, L. Turchi, R. Blanc-Mathieu, J. Lucas, P. Boddie, A. Khan, N. Manosalva Pérez, O. Fornes, T. Y. Leung, A. Aguirre, F. Hammal, D. Schmelter, D. Baranasic, B. Ballester, A. Sandelin, B. Lenhard, K. Vandepoele, W. W. Wasserman, F. Parcy, A. Mathelier, JASPAR 2022: the 9th release of the open-access database of transcription factor binding profiles. Nucleic Acids Res. 50, D165–D173 (2022).

