## Supplementary data for "FOXA2 is essential for maintaining the urea cycle in acute liver failure"

### SUPPLEMENTARY TABLES

**Table S1.** Immunohistochemical staining for CPS1, FOXA2 and C/EBPα in 20 ALF patients*

| | Patient | CPS1 | FOXA2 | C/EBPα |
|---|---|---|---|---|
| Non-HE | 1 | + | + | + |
| | 2 | + | + | + |
| | 3 | + | + | + |
| | 4 | + | + | + |
| | 5 | + | + | + |
| | 6 | + | + | + |
| | 7 | + | + | + |
| | 8 | + | + | - |
| | 9 | + | + | - |
| | 10 | + | + | - |
| | 11 | + | + | - |
| HE | 12 | - | + | - |
| | 13 | - | + | - |
| | 14 | - | + | - |
| | 15 | - | - | - |
| | 16 | - | - | - |
| | 17 | - | - | - |
| | 18 | - | - | - |
| | 19 | - | - | - |
| | 20 | - | - | - |

HE, hepatic encephalopathy

* Given the collected samples had been used in two previous studies [1,2], only 20 collected liver tissues had sufficient sample to perform all three IHC parameters.

**Table S2.** Characteristics of the study population

| | All (n=35) | Patients without HE (n=20) | Patients with HE (n=15) |
|---|---|---|---|
| **Age (years)** | 47.3 ± 9.7 | 48.2 ± 10.3 | 45.0 ± 8.2 |
| **Gender (F/M) (%)** | 14.3/85.7 | 13.3/86.7 | 25/75 |
| **Etiology of cirrhosis (%)** | | | |
| **-HBV** | 94.3 | 95 | 93.3 |
| **- Drug-induced liver injury** | 5.7 | 5 | 6.7 |
| **Ammonia (umol/L)** | - | - | 118 (58-310) |
| **Albumin (g/L)** | 30.6 ± 2.7 | 30.9 ± 3.3 | 30.5 ± 2.4 |
| **ALT (U/L)** | 575 (56-2019) | 417 (56-2019) | 607 (62-2019) |
| **AST (U/L)** | 205 (38-2437) | 159 (38-2437) | 348 (40-1854) |
| **Total bilirubin (umol/L)** | 305 (39-803) | 299 (39-726) | 316 (55-803) |
| **Creatinine (umol/L)** | 75 (43-188) | 71 (57-188) | 78 (43-168) |

HE, hepatic encephalopathy; HBV, hepatitis B virus; ALT, Alanine transaminase; AST, Aspartate aminotransferase

**Table S3.** Antibodies used in the study

| **IHC Staining** | | | | |
|---|---|---|---|---|
| Antibody | Species | Dilution | Company | Cat.No. |
| FOXA2 | Mouse | 1:100 | Santa Cruz | SC-101060 |
| CPS1 | Rabbit | 1:100 | Proteintech | 18703-1-AP |
| Anti-Rabbit/HRP | Swine | 1:200 | DAKO | P021702-2 |
| Anti-Mouse/HRP | Goat | 1:200 | DAKO | P0447 |

| **Immunoblotting** | | | |
|---|---|---|---|
| Antibody | Species | Company | Cat.No. |
| FOXA2 | Rabbit | Abcam | Ab108422 |
| CPS1 | Rabbit | Proteintech | 18703-1-AP |
| C/EBPα | Rabbit | Cell Signaling | CST#8178 |
| RARα | Rabbit | Cell Signaling | 62294S |
| GAPDH | Mouse | Santa Cruz | SC-25778 |
| Anti-Rabbit IgG | Mouse | Santa Cruz | SC-2357 |
| Anti-Mouse IgG | Goat | Santa Cruz | SC-2005 |

| **ChIP assay** | | | |
|---|---|---|---|
| Antibody | Species | Company | Cat.No. |
| RNA PoII Ser2 | Rabbit | Abcam | Ab193468 |
| RNA PoII Ser5 | Rabbit | Abcam | Ab5131 |
| H3K4me1 | Rabbit | Abcam | Ab8895 |
| H3K27ac | Rabbit | Abcam | Ab4729 |
| Histone3 | Rabbit | Abcam | Ab1791 |
| H3K4me3 | Rabbit | Abcam | Ab8580 |
| C/EBPα | Rabbit | Invitrogen | PA5-77911 |

| | | | |
|---|---|---|---|
| FOXA2 | Mouse | Sigma-Aldrich | 17-10258 |
| RARα | Rabbit | Cell Signaling | 62294S |
| IgG Control | Rabbit | Cell Signaling | CST#3900S |
| IgG Control | Mouse | Cell Signaling | CST#5415S |

**Table S4.** Primers used in the study

**ChIP-Primer**

| Gene | Forward primers (5’-3’) | Reverse primers (5’-3’) |
|---|---|---|
| *Cps1*(-718~-260bp) | AAACCGGCATATGGGAAGG | TGCACAGAGGGGAGTTGTTT |
| *Cps1*(-51~+80bp) | GGAGATAGAGGGGTTGGGTTT | CCTAAGCCACCCAGAGCATA |
| *Cps1*(-6222~-5644bp) | CCCATTCCAAATCTCACTGTGT | GGATGAGGACCTTGGAAACC |

**qPCR-Primer**

| Gene | Forward primers (5’-3’) | Reverse primers (5’-3’) |
|---|---|---|
| *Ppia* | GAGCTGTTTGCAGACAAAGTT | CCCTGGCACATGAATCCTGG |
| *Foxa2* | CCCTACGCCAACATGAACTCG | GTTCTGCCGGTAGAAAGGGA |
| *Cps1* | TACCCGGAAGCACTTACTGAT | GCCAGCCAGTGGTTATAGTCATT |
| *Otc* | AGGGTCACACTTCTGTGGTTC | CAGAGAGCCATAGCATGTACTG |
| *Cebpa* | GCAAAGCCAAGAAGTCGGTG | CACCTTCTGTTGCGTCTCCA |
| *Rara* | TCCGAAGAGATAGTACCCAGC | AGCCGGATGATTTGTCTTGAC |
| *Nags* | TACTCCTCCGCAGTCATTACA | CAGCATCCGCTCAGCATTTTT |
| *Arg1* | TGTCCCTAATGACAGCTCCTT | GCATCCACCCAAATGACACAT |
| *Ass1* | GCCTTGCATAGCTCGCAGA | GGACCTGGTCATTCCCCTTT |
| *Asl* | CAGGACGAAGTCGCAACGA | GGAGGGCGGAGAGTTTTGAG |
| *GAPDH* | GGAGCGAGATCCCTCCAAAAT | GGCTGTTGTCATACTTCTCATGG |
| *FOXA2* | CCAGGAATTTGCTGCTCTTC | TCCATAGGGACCACACACAA |
| *CPS1* | AATGAGGTGGGCTTAAAGCAAG | AGTTCCACTCCACAGTTCAGA |
| *OTC* | CAGAGGCAGAAAACAGAAAGTGG | ACAATGGCAAAGCATATCATACGG |
| *CEBPA* | TATAGGCTGGGCTTCCCCTT | AGCTTTCTGGTGTGACTCGG |
| *RARA* | GCTAAAGGTCTACGTGCGGA | GGCCCTCTGAGTTCTCCAAC |
| *NAGS* | TGAGTAACGTGAACCTGCCC | CCGGACCCCTTGTTGCTAAA |
| *ARG1* | AAGATTCCCGATGTGCCAGG | GTCCACGTCTCTCAAGCCAA |
| *ASS1* | CGGGAAGTGCGCAAAATCAA | CACTTTCCCTTCCACTCGCT |

| | | |
|---|---|---|
| *ASL* | ACGCTGCAGATTCACCAAGA | CGGCCATGAACACAGCTTTC |

**Table S5.** siRNA and plasmid used in the study

| **Small interfering RNA** | | | |
|---|---|---|---|
| Product | Species | Company | Cat.No. |
| siCEBPA | Human | Santa Cruz | SC-37047 |
| siCebpa | Mouse | Santa Cruz | SC-37048 |
| siFOXA2 | Human | Santa Cruz | SC-35569 |
| siFoxa2 | Mouse | Santa Cruz | SC-35570 |
| siRARA | Human | Santa Cruz | SC-29465 |
| siRara | Mouse | Santa Cruz | SC-36393 |
| Control siRNA | | Santa Cruz | SC-37007 |
| **Recombinant DNA** | | Company | Cat.No. |
| pFlag-CMV4-FOXA2 | | Biomed | BM1305 |
| pFlag-CMV4-CEBPA | | Biomed | BM1310 |
| pFlag-CMV4 | | Biomed | |
| pLV.PGK.mFoxa2 | | Addgene | 33014 |
| pScalps-mcebpa | | Addgene | 79551 |
| pGCDNsam-Hnf4a-IRES-GFP | | Addgene | 33006 |

**Table S6. The sequences of siRNA**

| siRNA | **Sense (5'-3')** | **Antisense (5'-3')** |
|---|---|---|
| siCEBPA (sc-37047A) | CUCCAAUGCCUACUGAGUAtt | UACUCAGUAGGCAUUGGAGtt |
| siCEBPA (sc-37047B) | CCACGCCUGUCCUUAGAAAtt | UUUCUAAGGACAGGCGUGGtt |
| siCEBPA (sc-37047C) | CCGAUAUCAACACUUGUAUtt | AUACAAGUGUUGAUAUCGGtt |
| siCebpa (sc-37048A) | CGCUGGUGAUCAAACAAGAtt | UCUUGUUUGAUCACCAGCGtt |
| siCebpa (sc-37048B) | CCCACUUGCAGUUCCAGAUtt | AUCUGGAACUGCAAGUGGGtt |
| siCebpa (sc-37048C) | AGACCACCAUGCACCUACAtt | UGUAGGUGCAUGGUGGUCUtt |
| siFOXA2 (sc-35569A) | CCCAUUAUGAACUCCUCUUtt | AAGAGGAGUUCAUAAUGGGtt |
| siFOXA2 (sc-35569B) | GACAAGUGAGAGAGCAAGUtt | ACUUGCUCUCUCACUUGUCtt |
| siFOXA2 (sc-35569C) | GUUCUCCUCCAUUGCUGUUtt | AACAGCAAUGGAGGAGAACtt |
| siFoxa2 (sc-35570A) | CACCCCUUCUCUAUCAACAtt | UGUUGAUAGAGAAGGGGUGtt |
| siFoxa2 (sc-35570B) | GUAGACUACUGCUUCUCAAtt | UUGAGAAGCAGUAGUCUACtt |
| siFoxa2 (sc-35570C) | GGGUUGUACUGAUGUUGAAtt | UUCAACAUCAGUACAACCCtt |
| siRARA (sc-29465A) | CAGCAGUUCUGAAGAGAUAtt | UAUCUCUUCAGAACUGCUGtt |
| siRARA (sc-29465B) | CCAAGGAGUCUGUGAGAAAtt | UUUCUCACAGACUCCUUGGtt |
| siRARA (sc-29465C) | GAGAUCUACUGGAUAAAGAtt | UCUUUAUCCAGUAGAUCUCtt |
| siRara (sc-36393A) | GCCUUGCUUUGUUUGUCAAtt | UUGACAAACAAAGCAAGGCtt |
| siRara (sc-36393B) | CGAGCUCAUUGAGAAGGUUtt | AACCUUCUCAAUGAGCUCGtt |
| siRara (sc-36393C) | GGCAAGUACACUACGAACAtt | UGUUCGUAGUGUACUUGCCtt |

**Table S7.** Reagent kits used in the study

| **Commercial assays** | | |
|---|---|---|
| Product | Company | Cat.No. |
| Bio-Rad protein assay kit | Bio Rad | 5000113-115 |
| Lipofectamine® RNAiMAX kit | Thermo Fischer | 13778-075 |
| Lipofectamine 3000 kit | Thermo Fischer | L3000008 |
| PureLink Quick Plasmid Miniprep Kit | Invitrogen | K210010 |
| PureLink® HiPure Plasmid Maxiprep Kit | Invitrogen | K210007 |
| Urea Assay Kit | Abcam | Ab83362 |
| Ammonia Assay Kit | Abcam | Ab83360 |
| Phusion® High-Fidelity DNA Polymerase | New England Biolabs | M0530S |
| SYBR Green Master Kit | Thermo Fisher | A25918 |
| ALT/AST KIT | Shenzhen Mindray Bio-Medical Electronics Co. | 105-000442-00<br>105-000443-00 |

### SUPPLEMENTARY FIGURES AND FIGURE LEGENDS

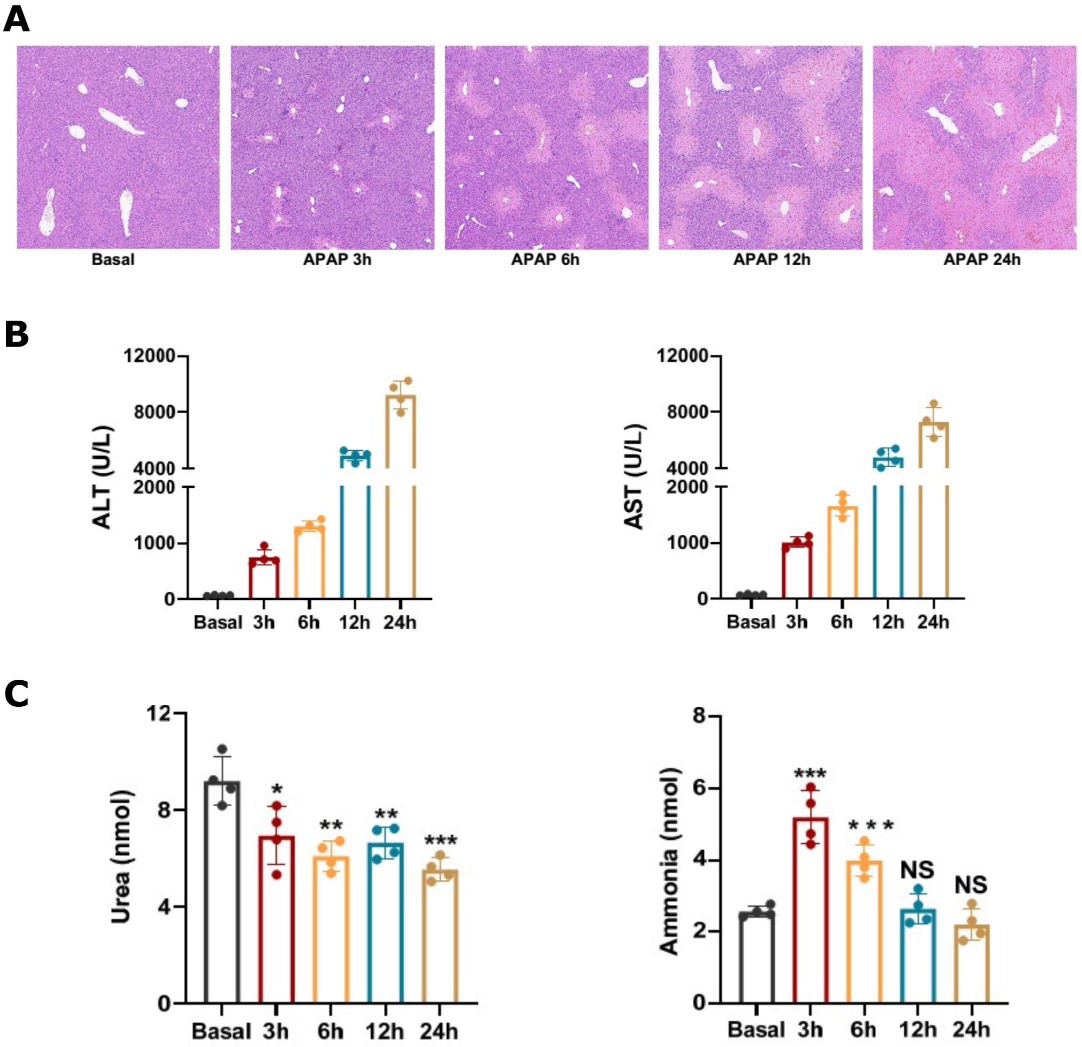


**Fig. S1. Acetaminophen injections induce liver injury in mice at different time points**

**(A).** H&E staining showed the necrotic areas in the live tissues of mice with indicated administration. **(B)** ALT, AST, **(C)** urea and ammonia concentrations in serum samples were collected at different time points after APAP injection and were analyzed as described in Materials and Methods section. The samples were collected at 3h, 6h, 12h and 24h following APAP injection. Bars represent the mean ± SD, *P* values were calculated by One-way ANOVA, *, $P<0.05$; **, $P<0.01$, and ***, $P<0.001$ and NS, no significance.

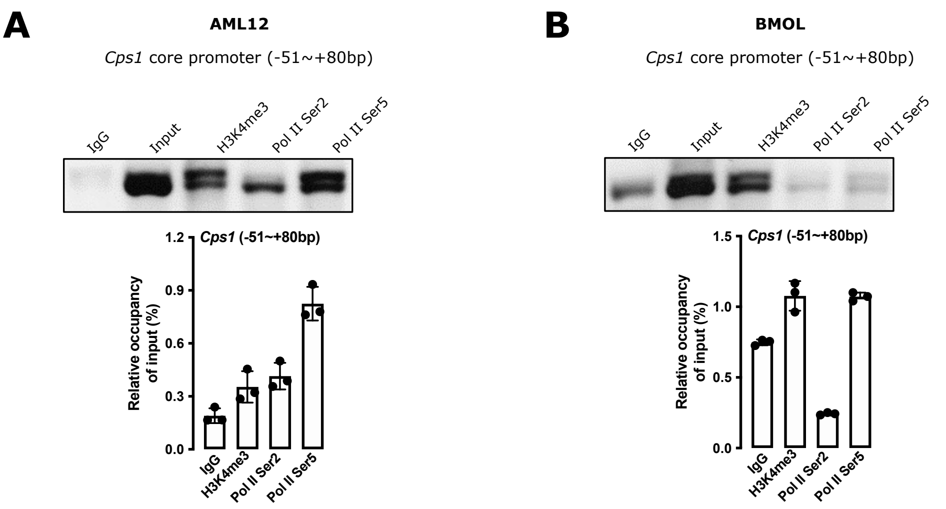


**Fig. S2. The analysis of *Cps1* core promoters in hepatocytes and liver progenitor cells**

**(A-B).** ChIP assays were performed to examine H3K4me3, Pol II S2 and Pol II S5 binding to *Cps1* (-51~+80bp) core promoters in AML12 **(B)** and BMOL cells **(C)**. The binding activities were analyzed by ChIP-qPCR. Bars represent the mean ± SD. Triple experiments were performed, and one representative experiment is shown.

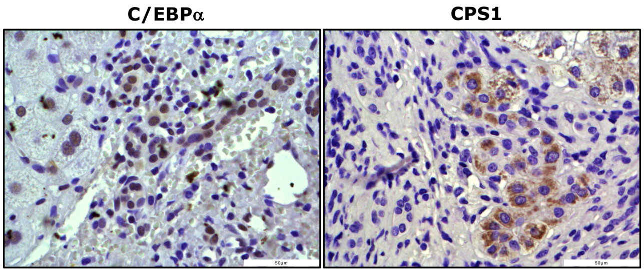


**Fig. S3. C/EBPα and CPS1 expression in cirrhotic patients with ductular reactions**

Immunohistochemical staining for C/EBPα and CPS1 was performed in cirrhotic patients with ductular reactions. Representative patients show that C/EBPα and CPS1 are expressed in liver progenitor cells (LPC) in cirrhotic patients with ductular reactions. Original magnification: x40.

### REFERENCES

1. Feng R, Kan K, Sticht C, et al. A hierarchical regulatory network ensures stable albumin transcription under various pathophysiological conditions. *Hepatology*. Published online February 19, 2022. doi:10.1002/hep.32414
2. Feng R, Tong C, Lin T, et al. Insulin Determines Transforming Growth Factor β Effects on Hepatocyte Nuclear Factor 4α Transcription in Hepatocytes. *Am J Pathol*. Published online October 10, 2023:S0002-9440(23)00365-6. doi:10.1016/j.ajpath.2023.09.009
